# Deactivation of SARS-CoV-2 surrogate porcine epidemic diarrhea virus with electron beam irradiation under the cold chain transportation condition

**DOI:** 10.1101/2021.09.25.461766

**Authors:** Yan Liu, Yang Shao, Lu Wang, Weilai Lu, Shihua Li, Diandou Xu, Yu Vincent Fu

## Abstract

Severe acute respiratory syndrome coronavirus 2 (SARS-CoV-2) has prevailed all over the world and emerged as a significant public health emergency. The rapid outbreak of SARS-CoV-2 is largely due to its high transmission capacity. Studies implied that the cold chain logistics would be a potential route for the spread of SARS-CoV-2. The low temperature condition of the cold chain is conducive to survival and transmission of virus. Thus, the virus disinfection in cold chain should not be neglected for controlling COVID-19. However, due to the low temperature feature of the cold-chain, the virus disinfecting methods suitable in cold chain are limited. Here the high-energy electron beam irradiation is proposed to disinfect the SARS-CoV-2 in cold chain logistics. We evaluated the impact of high-energy electron beam irradiation on porcine epidemic diarrhea virus (PEDV), an enveloped virus surrogate for SARS-CoV-2, and explored the possible mechanism of the action of high-energy electron beam irradiation on PEDV. The irradiation dose of 10 kGy inactivated 98.1 % PEDV on the both top and bottom surfaces of various packaging materials under cold chain frozen condition. High-energy electron beam inactivated PEDV by inducing damages on viral genome or even capsid.

## 1. Introduction

Viruses are submicroscopic infectious agents that cause many serious diseases in human. To help prevent the infectious diseases caused by viruses, one of the important ways is deactivating virus in its route of transmission out of human body. The recent COVID-19 pandemic caused by SARS-CoV-2 brought a massive disaster to human beings globally. SARS-CoV-2 belongs to coronavirus (CoV), which is an enveloped, single-stranded, positive-sense RNA virus. Many studies have demonstrated that temperature is the most influential factor for the survivability of CoV in inanimate environments, and lower temperature leads to a slow viral inactivation(1). SARS-CoV-2 has been reported to remain viable up to at least 14 days at 4 °C, while the inactivation time is 5 minutes at 70 °C (2, 3). Riddell et al. showed that infectious SARS-CoV-2 virus were recovered after at least 28-day incubation at 20 °C (4). Some studies demonstrated that CoV persisted infectiously to several months or up to a year at 4 °C and lost little infectivity for many years when keep at -60 °C(5, 6). It has been reported that infectious bovine coronavirus were detected on lettuce surface after at least 14-day storage at 4 °C, and that a wash procedure did not remove residual viruses completely (7). Therefore, the contaminated foods might be potential vehicles for coronavirus infection, especially for the food transported via cold chain, since CoVs are stable at low temperature.

Recently, it has reported that surface swab samples related to imported cold chain food were tested SARS-CoV-2 nucleic acid positive in Beijing, Tianjin, Dalian and Qingdao, China (8, 9). Notably, the viable SARS-CoV-2 were isolated from the cold chain outer package surface of imported cod in Qingdao (http://www.chinacdc.cn/). As suggested by Ji et al., the virus test and disinfection on cold chain should not be neglected for controlling COVID-19(8). However, virus disinfection in cold chain logistics is not an easy job. So far, heating, chemical disinfection and UV irradiation are the three most common-used virus disinfecting methods. Heating is obviously not suitable for cold-chain. Water-dissolved chemical disinfection may not be applied in the cold chain logistics due to the low temperature (normally below zero) must be maintained. In addition, the efficiency of chemical disinfection could be dramatically affected by the low temperature of cold chain. It is known that UV irradiation can deactivate SARS-CoV-2 at room temperature (10, 11). However, the penetration of UV irradiation is limited especially under the frozen situation in cold chain.

Irradiation technology is a microbial inactivating method suitable for cold chain because it is non-thermal technique with high penetration and leaving no residue after the processing. Among different types of irradiation, E-beam irradiation, which uses high voltage power supply and accelerator to produce high energy electrons, is an emerging technology to catch more interests. Compared to gamma irradiation, E-beam irradiation is more environment-friendly. Moreover, for products that are sensitive to oxidative effects, E-beam irradiation is safer than gamma exposure. Some studies also showed that the treatment time of E-beam irradiation is generally shorter than gamma irradiation because of the higher dose rates of E-beam irradiation(12-16). E-beam irradiation has been reported to efficiently inactivate Tulane virus, murine norovirus, influenza A (H3N8), porcine reproductive and respiratory syndrome virus and equine herpesvirus 1(17, 18). So far, no studies examined the efficacy of E-beam irradiation on coronavirus under the cold chain logistics condition.

In order to examine whether E-beam irradiation can efficiently deactivate SARS-CoV-2 in cold-chain condition, we evaluated the sensitivity of SARS-CoV-2 surrogate, porcine epidemic diarrhea virus (PEDV), to E-beam irradiation on different surface in frozen condition. Our data showed that the irradiation dose of 10 kGy effectively inactivated PEDV on the both top and bottom surfaces of common used wrapped materials in frozen condition. After E-beam irradiation treatment, a massive RNA breakage was observed on the PEDV genome, plus the capsid damages under a high dose irradiation, suggesting a mechanism for E-beam to inactivate CoV.

## 2. Materials and Methods

### 2.1. Virus propagation and cell culture

Vero E6 cell was a gift from Dr. George F. Gao in the Institute of Microbiology, Chinese Academy of Sciences. Cells were maintained in Dulbecco’s modified Eagle medium (DMEM) supplemented with 10% fetal bovine sera (FBS), 200 mg/ml streptomycin, 200 IU/ml penicillin and cultured at 37°C with 5% CO_2_. Porcine Epidemic Diarrhea virus (PEDV) strain PEDV-HB3 was a gift provided by Dr. Jinghua Yan in the Institute of Microbiology, Chinese Academy of Sciences. All the viral experiments were performed in a biosafety level-2 (BLS-2) laboratory.

For the PEDV-HB3 virus propagation assay, 100 μL PEDV-HB3 virus solution was add to the overnight cultured Vero E6 cells and incubated at 37 °C with 5% CO_2_ for 1 h to allow virus particle to attach to the Vero E6 cells. After removing the media (DMEM containing unabsorbed viruses) and washing the cells with PBS, 20 ml cell culture maintenance medium (DMEM containing 7ng/μL Trypsin) was add to the cells and incubated for 24-48 h to propagate the virus. We performed three cycles (alternating cycles of -80 °C and 25 °C) of freezing and thawing to separate the virus from the Vero E6 cells(19). Cellular debris was removed by centrifugation at 800g for 5 min and then filtered through a 0.22 μM filter. The harvested virus was aliquoted and stored at -80°C until use. Then the virus titers were determined by TCID_50_ assay described in Section 2.4.

### 2.2. Stability of PEDV on the surfaces at -20 °C

Surface stability was assessed on corrugated cardboard, polystyrene plastics and polystyrene foam representing a variety of cold-chain transport materials and was carried out as described by Neeltje et al. (2020) with slight modifications(20). In short, 50 μL of PEDV with a titer of 10^4.15^ TCID_50_/100μL was deposited on the surface. After being placed at -20°C for 7 days, 14 days, and 30 days, 1 mL of DMEM was added to recover the inoculated virus. Virus titer was immediately measure by TCID_50_ assay described in Section 2.4. All experiments were performed in at least triple repeats.

### 2.3. Inactivation of PEDV by E-beam irradiation

PEDV-HB3 with a titer of 100 TCID_50_/100μL was deposited on the top or bottom surfaces of three cold-chain transport materials (corrugated cardboard, polystyrene plastics and polystyrene foam). The average thickness of corrugated cardboard, polystyrene plastics and polystyrene foam used in the experiments is 4.9 mm, 0.1 mm, 18.2 mm respectively. The samples were frozen under -20 °C and irradiated with the predefined doses of 0, 2, 5, 10, 20, 25, and 30 kGy on dry ice with a 10 MeV electron accelerator. The E-beam irradiated viruses were stored at -80 °C until subsequent analyses.

All experiments were performed in at least triple repeats.

### 2.4. Viral infectivity detection by TCID_50_ assay

The virus titer was determined using 50% tissue culture infective dose (TCID_50_) assay(21). Briefly, one hundred microliters of tenfold serially diluted virus stock (each dilution had eight replicates) infected overnight cultured Vero E6 cells in a 96-well plate for 72 h. After incubation, the plate was examined for the virus-induced cell cytopathic effect (CPE) in each well via microscopy. Virus titers were calculated using the Karber method(22). Similarly, viable virus counts after E-beam irradiation were also determined by TCID_50_ assay. Specifically, when the post-E-beam irradiated viruses did not cause CPE, blind transmission of three generations was performed as follows to determine that the irradiated virus had no infective activity. After observing CPE, the virus was harvested and inoculated into the cells for infection as in Section 2.1(repeat this procedure 2 times). In the meanwhile, once CPE was present, the blind transmission experiment was stopped.

### 2.5. Viral capsid damage was determined by RNase assay

We conducted a ribonuclease (RNase) assay that was modified from Chamteut et al.(2020) and Elbashir et al.(2020) to evaluate the integrity of the PEDV-HB3 capsid(23, 24). For this assay, a 140-μL volume of unirradiated or irradiated PEDV-HB3 was add to a 1.5-mL RNase-free microcentrifuge tube. Next, 2.8 μL of RNase (1mg/ml) was added to each tube. After the mixture was incubated in a water bath at 37 °C for 30 min, 1 μL of RNase inhibitor(40 U/μL) was added to the mixture and incubated at room temperature for another 30 min. Here, we used the RNase inhibitor to remove the remaining RNase from the reaction. Correspondingly, 140 μL of virus before RNase treatment was used as control. Viral capsid damage was determined by the reduction in viral RNA copy number caused by RNase degradation. RNA was extracted from 140 μL of PEDV-HB3 solution using a QIAamp Viral RNA mini kit (QIAGEN), following the manufacturer’s instructions. The extracted RNA was quantified by a HiScript II U+ One Step qRT-PCR Probe Kit (Vazyme) according to the manufacturer’s instructions. The primers and the TaqMan probe targeting N gene of PEDV-HB3 were listed in Table 1. Meantime, a PEDV-HB3 standard curve was used to quantify the RNA copy number (Table 1). Copy numbers of each RNase-treated PEDV-HB3 sample was divided by that of control PEDV-HB3(samples were not treated with RNase) to calculate the copy number ratio. Assay was performed in a triplicate experiment.

**Table 1.**
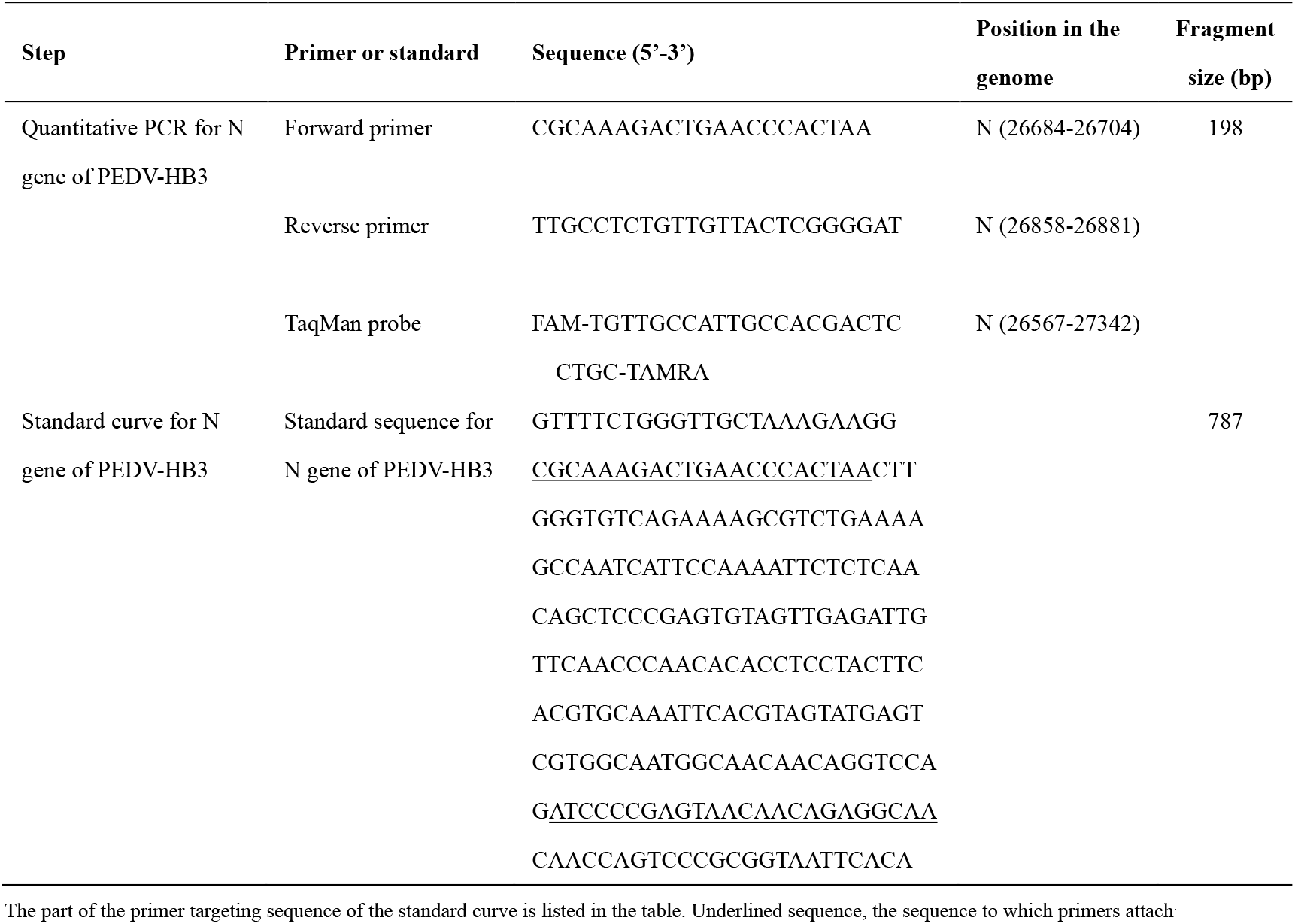
Primers and standard for detecting N gene of PEDV-HB3 by one-step RT-qPCR

### 2.6. Virus genomic damage was determined by long-range RT-qPCR assay

We conducted a long-rang RT-qPCR assay to analyze the effect of E-beam irradiation on the viral genome. RNA was extracted from 140 μL of virus samples irradiated with different doses. Then, we used SuperScript^®^ III First-Strand Synthesis System for RT-PCR (Invitrogen) to perform reverse transcription on 3 μL volume of RNA according to the standard protocol of the manual. This long-rang reverse transcription with specific reverse transcription primer located at 21594 bp of the PEDV-HB3 genome was carried out to ensure at least 8 K cDNA products from viral genome (Table 2). Three qPCR primer sets were designed at 746 bp, 3655 bp and 7674 bp upstream of the reverse transcription primer site, respectively (Table 2). Viral cDNA was then subjected to qPCR using Hieff® qPCR SYBR® Green Master Mix (Yeasen), following the manufacturer’s instructions. A PEDV-HB3 standard curve was used to quantify the RNA copy number. Here, PMD18-T plasmid containing S gene fragment was constructed as the standard (Table 2). The viral genomic damage rate was calculated as follows:

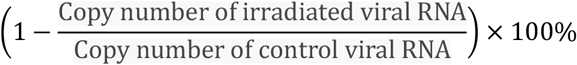

All experiments were performed in at least triple repeats.

**Table 2.**
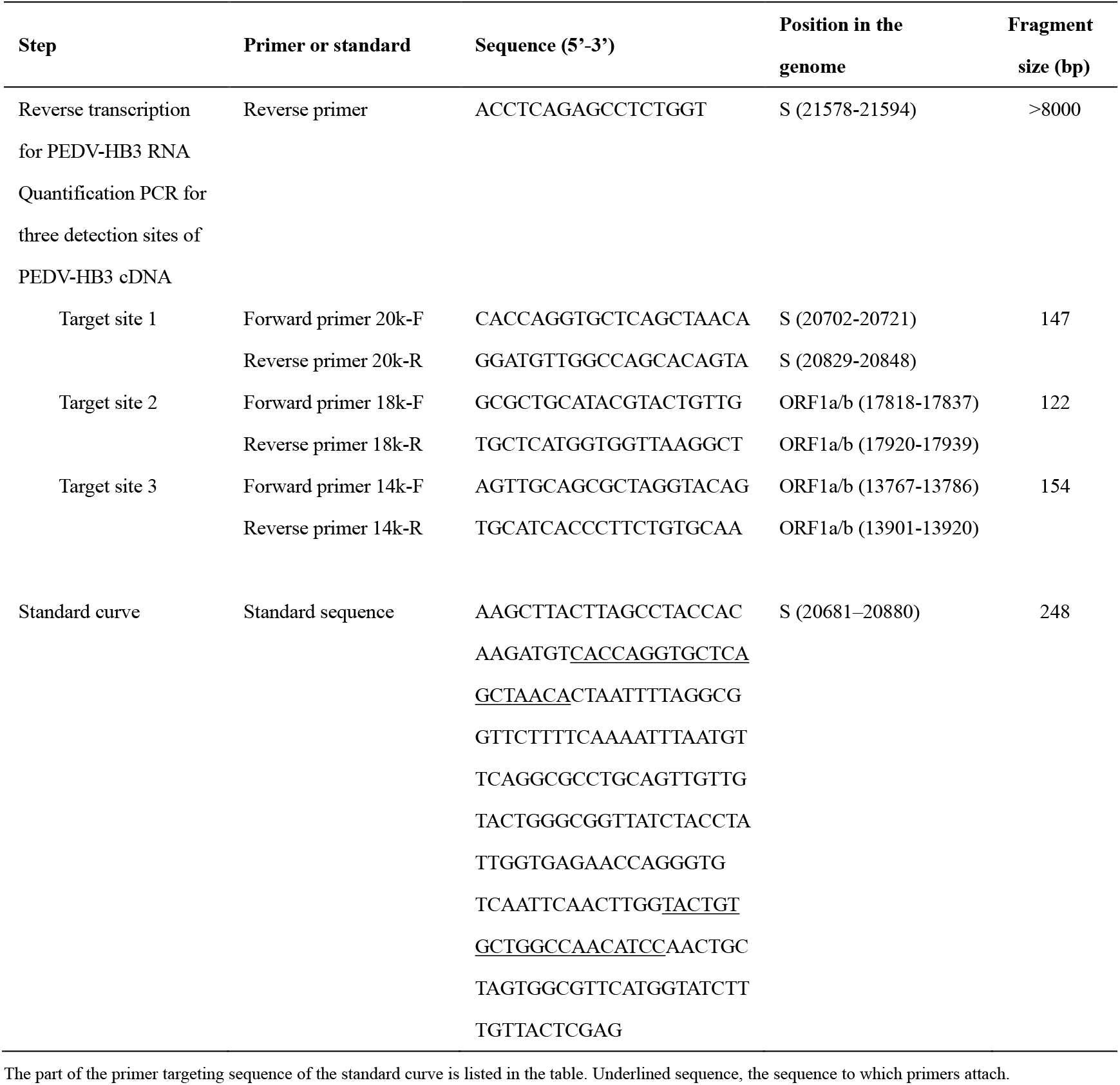
Primers and standards for detecting genome damage by two-step RT-qPCR

### 2.7. Statistical analysis

All assays were performed in at least triple replicates. Student’s t-test was used to evaluate the significance between datasets of TCID_50_ assay. One-way analysis of variance (ANOVA) followed by Dunnett’s multiple comparisons test was used to determine the significance in RT-qPCR. P value less than 0.05 indicates a significant difference between the two datasets.

## 3. Results

### 3.1. PEDV-HB3 remained infectious activity at least 30 days under -20 °C

To assess the stability of coronavirus under cold chain conditions, PEDV-HB3 was inoculated on the surface of three common cold chain packaging materials (corrugated cardboard, polystyrene plastics and polystyrene foam) with an inoculum of 10^4.15^ TCID_50_/100μL and kept at -20 °C. We collected samples at specified time-points and analyzed the infectious activity of virus by using TCID_50_ assay. As shown in Figure 1, the PEDV remained viable on the package surfaces throughout the duration of 30 days at -20 °C. The PEDV infectious titer only reduced 2 to 3 logs after 30 days (from 4.15 lgTCID_50_/100μL to 1.58 lgTCID_50_/100μL after 30 days on corrugated cardboard, from 4.15 lgTCID_50_/100μL to 2.08 lgTCID_50_/100μL after 30 days on polystyrene plastics and from 4.15 lgTCID_50_/100μL to 1.54 lgTCID_50_/100μL after 30 days on polystyrene foam). Virus on the various surfaces showed different stability. The infectious titer of virus on the porous material (corrugated cardboard and polystyrene foam) surface was reduced more greatly than that on the non-porous materials (polystyrene plastics) at all time-points. Despite the effects on virus viability from the chemical and physical characters of different packaging materials, it is possible that the observed difference was partially due to the recovery efficiency of virus from different surfaces.

**Figure 1.**
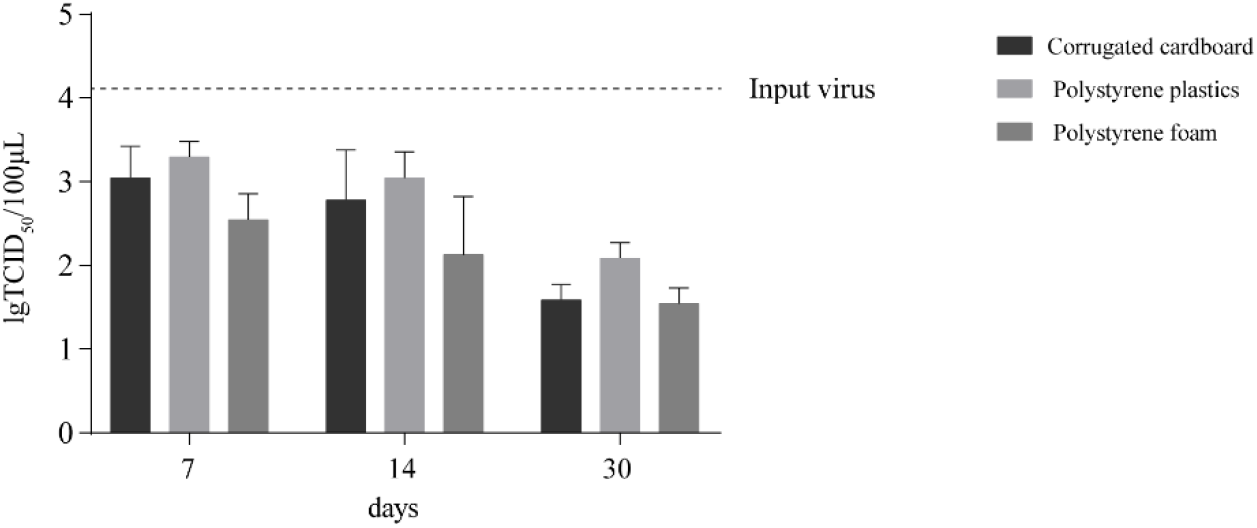
Virus stability on the surfaces of three common cold chain packaging materials at -20 °C. Virus was recovered at indicated time-points and titrated for analyzing infectious activity. Error bars represent the standard deviation (SD) of triple replicates.

### 3.2. E-beam inactivated PEDV-HB3 in a dose-dependent manner

To test the inactivation efficiency of E-beam irradiation on coronavirus under the cold chain condition, PEDV-HB3 deposited on three common cold chain packaging materials (corrugated cardboard, polystyrene plastics and polystyrene foam) at both top and bottom surfaces were irradiated with the predefined doses of E-beam at -20 °C. TCID_50_ assay was then carried out to quantify the virus infectious titer. A marked cytopathic effect was produced in cell cultures by the PEDV without E-beam irradiation (Figure 2A, see “0 kGy”), and the infectious titer was shown in Table 3. In contrast, no cytopathic effects were observed in the cell cultures infected by the PEDV dealt with 10 kGy, 20 kGy, 25 kGy, and 30 kGy radiation dose. These cell cultures showed the normal morphology same as the cells without virus infection treatment (Figure 2A, see “10 kGy, 20 kGy, 25 kGy and 30 kGy “). As shown in Table 3, the infectious titers of these irradiated virus were ≤-0.5 lgTCID_50_/100μL. After treating with the PEDV irradiated with 2 kGy and 5 kGy dose, a small number of cells displayed marked cytopathic effects (Figure 2A, see “2 kGy and 5 kGy “). The corresponding virus infectious titers were shown in Table 3. After the PEDV dealt with 2 kGy radiation dose, the infectious titer reduced 96.3%. The reduction of infectious titer was 96.9% with a radiation dose of 5 kGy (Table 3). No significant difference in the reduction ratio of infectious tires was observed for the PEDV on the top surfaces of different packaging materials (Figure 2C). Similarly, there was no significant difference in the infectious titer reduction for the PEDV deposited on the top surface or bottom surface of a packaging material (Figure 2C). These data suggested that a rather low dose E-beam irradiation (10 kGy) can largely inactivate PEDV-HB3 with an at least 18.2 mm penetrating distance for polystyrene foam, 4.9 mm for corrugated cardboard and 0.1 mm for polystyrene plastics under cold chain frozen condition.

**Figure 2.**
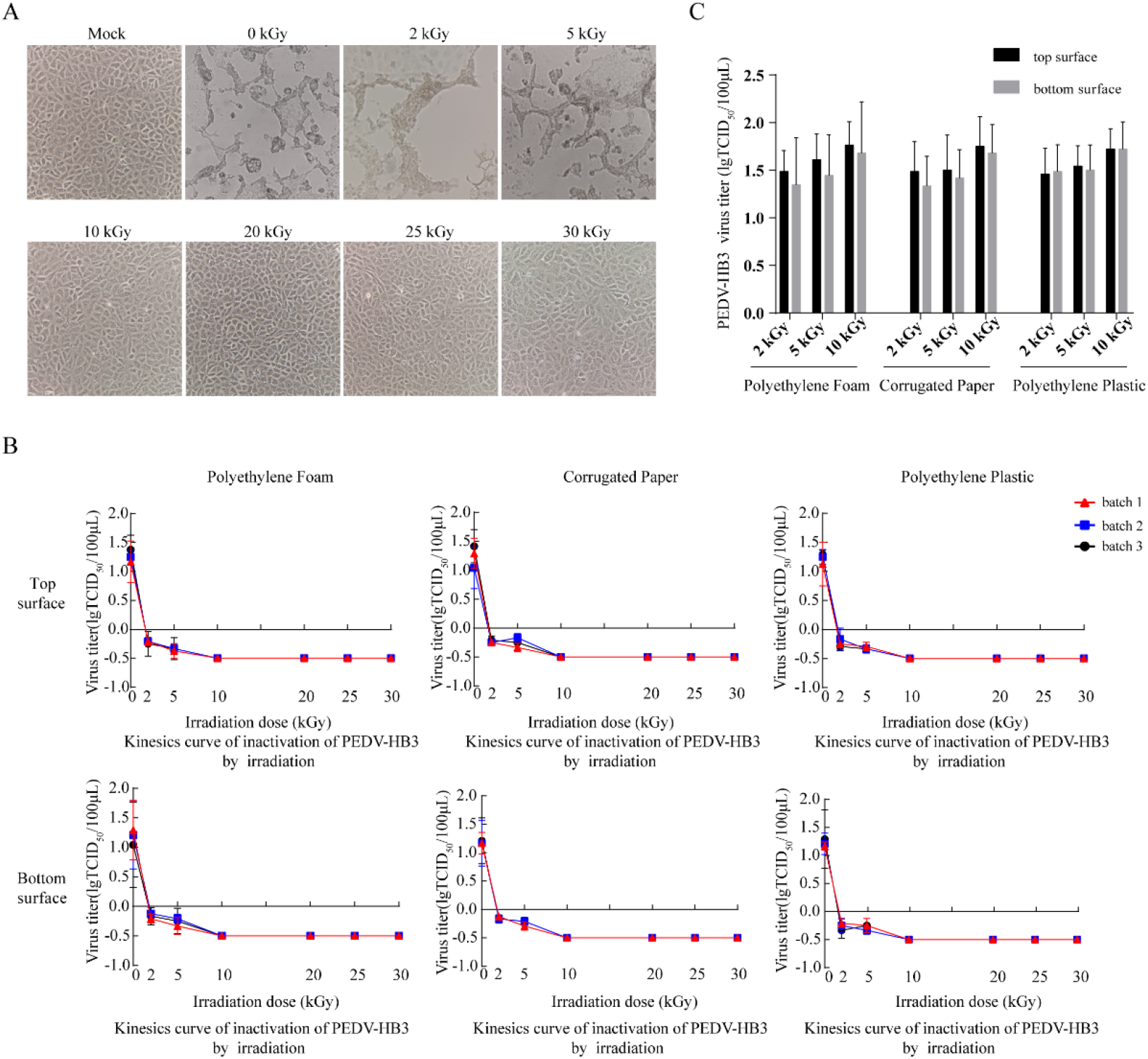
Deactivating effects of E-beam-irradiation on PEDV-HB3. (A) Cytopathic changes in Vero E6 cells inoculated with the PEDV-HB3 without E-beam irradiation (0 kGy), or with E-beam irradiation with 2, 5, 10, 20, 25 or 30 kGy. The representative images of cell cultures are shown. (B) Kinesics curve of inactivation of PEDV-HB3 by E-beam irradiation. The mean of infectious titers were plotted. Error bars represent the standard deviation (SD) of triple replicate experiments. (C) Analysis of the significant difference of infectious titers on various materials or on the top and bottom surfaces after predefined dose E-beam irradiation. Student’s t-test was used to evaluate the significance between datasets of TCID_50_ assay. P value less than 0.05 indicates a significant difference between the two datasets.

**Table 3.**
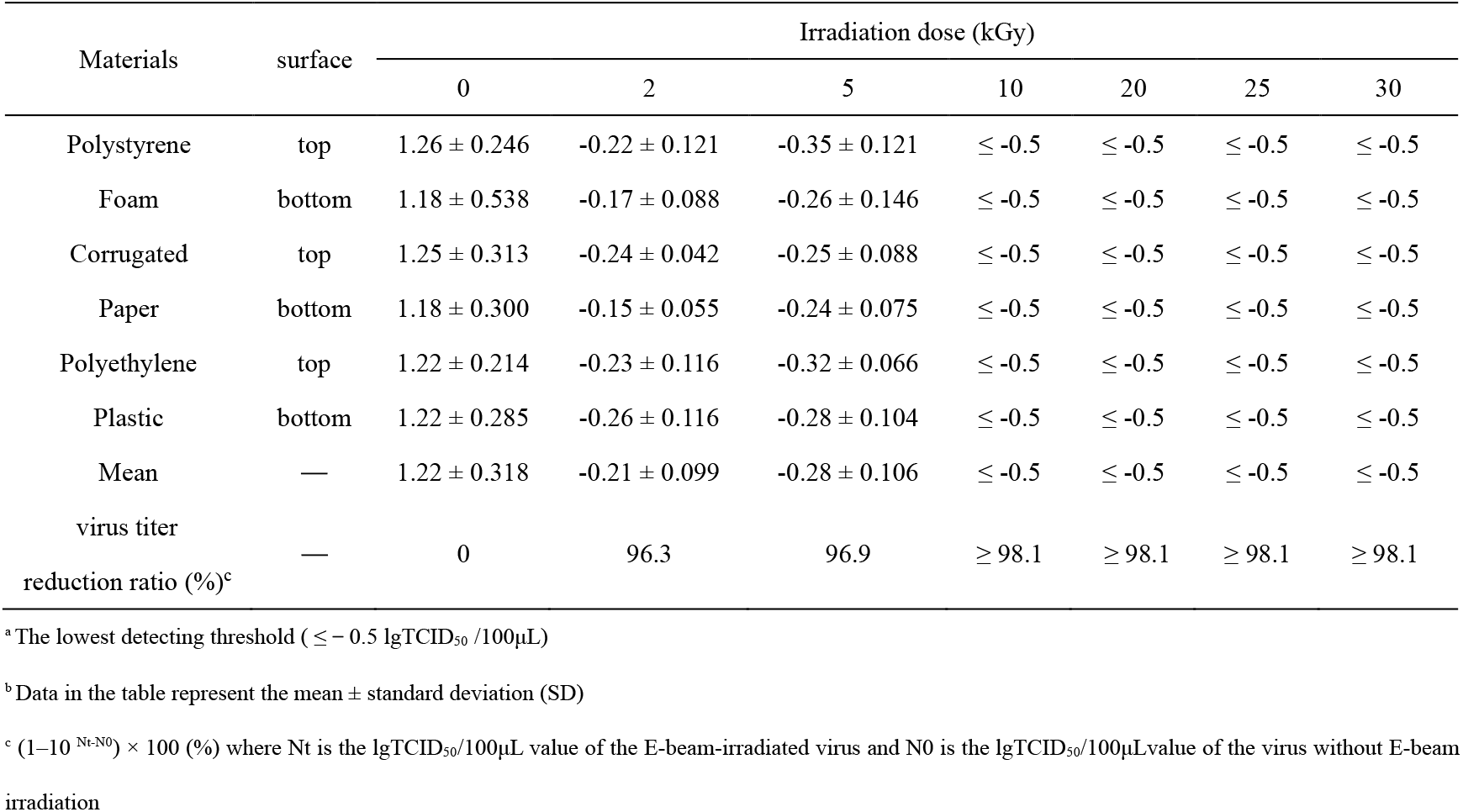
PEDV-HB3 virus titer (lgTCID_50_/100μL) after irradiation at predefined dose

The kinesics curve of PEDV-HB3 inactivation by E-beam irradiation analysis showed that E-beam inactivated PEDV-HB3 in a dose-dependent manner. The E-beam irradiation of 10 kGy dose completely abolished PEDV-HB3 infectivity (Figure 2B). According to the requirements of “National Drug Administration Note [2002] No. 160” of China, if the virus titer reduction of less than 4 lgTCID_50_/100μL, the inactivated virus samples should be blindly transmitted for 3 generations to ensure the inactivation. Therefore, virus irradiated with 10, 20, 25 and 30 kGy dose were blindly transmitted for three generations. As shown in Table 4, cytopathic effects were not observed in all cell cultures on each generation. Combining all data together, a 10 kGy radiation dose can effectively deactivate PEDV-HB3 on the both top and bottom surfaces of common used packaging materials in cold chain under the – 20 °C condition.

**Table 4.** The induced cytopathy in cells of three-generation blind transmission after PEDV inoculation

| Materials | surface | Irradiation dose (kGy) |  |  |  |  |
| --- | --- | --- | --- | --- | --- | --- |
|  |  | 0 | 10 | 20 | 25 | 30 |
| Polystyrene | top | + | — | — | — | — |
| Foam | bottom | + | — | — | — | — |
| Corrugated | top | + | — | — | — | — |
| Paper | bottom | + | — | — | — | — |
| Polyethylene | top | + | — | — | — | — |
| Plastic | bottom | + | — | — | — | — |

cytopathogenic effect, CPE: +

### 3.3. High dose E-beam irradiation disrupted the integrity of PEDV-HB3 capsid

In order to decipher the mechanism how the E-beam inactivates the PEDV, we tested whether E-beam directly disrupts the integrity of viral capsid using an RNase assay. An intact capsid can efficiently prevent the access of RNase to viral RNA, thus RNase prefers to degrade the viral RNA when the viral capsid is disrupted. The damage of viral capsid was determined based on the reduction of viral RNA copy number caused by RNase degradation. Briefly, PEDV-HB3 was treated with E-beam irradiation. After treatment, RNase was used to degrade RNA in the disrupted virions, followed by RT-qPCR to quantify the viral RNA copy number. PEDV was deposited on the top surface of polystyrene plastics and treated with predefined dose of E-beam irradiation under -20 °C. When treating the virus with low radiation dose (0 kGy, 5 kGy and 10 kGy), the reduction of viral RNA copy number showed no significant differences between the non-treatment control group and low dose irradiated group (p < 0.05) (Figure 3). However, if the radiation dose reached to 20 kGy, 25 kGy or 30 kGy, the reduction in viral RNA copy number was significantly higher than the non-treatment control (Figure 3). It suggested that the disruption of PEDV capsid required a high dose E-beam irradiation. E-beam irradiation of 10 kGy does not cause obvious damage to the integrity of PEDV-HB3, meanwhile, 10 kGy E-beam irradiation can efficiently inactivate PEDV. These data implied the major mechanism of E-beam-induced PEDV inactivation with irradiation might not due to the damage to viral capsid.

**Figure 3.**
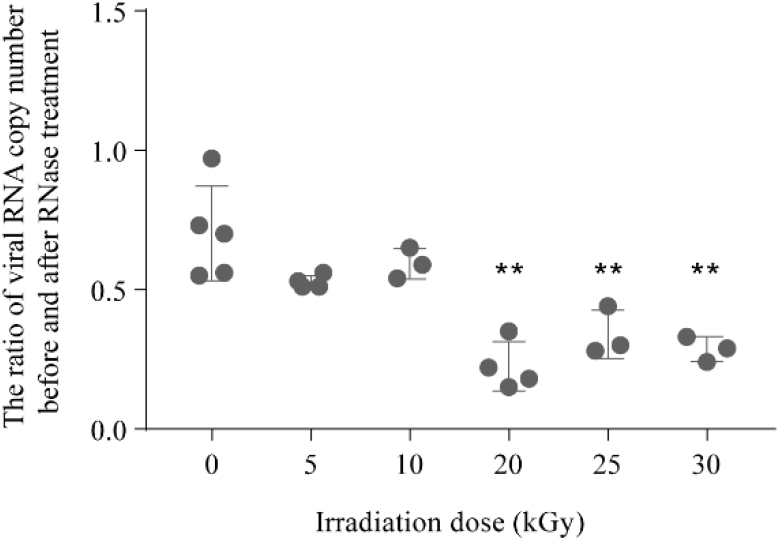
The effect of E-beam irradiation on the PEDV-HB3 capsid. The integrity of viral capsid was determined by the reduction of viral RNA copy number which was caused by RNase treatment. The reduction ratio of viral RNA copy number was defined as the ratio of viral RNA copy number before and after RNase treatment. Experiments of indicated dose (0 kGy, 5 kGy, 10 kGy, 20 kGy, 25 kGy and 30 kGy) were repeated at least three times as described in Methods and Materials. Each solid point represents the result from an independent experiment. Significance was determined using a one-way analysis of variance (ANOVA) followed by Dunnett’s multiple comparisons test (p < 0.05). Error bars represent the standard deviation (SD) of replicate experiments.

### 3.4. E-beam irradiation damaged the PEDV genome

Previous studies have shown that the damage on viral RNA genome is strongly correlated with virus infectivity (24). E-beam is the machine-accelerated electrons so that it has the ability to break viral RNA strand for inactivating PEDV. Therefore, the mechanism of virus disinfection by E-beam could be the breakages on viral RNA genome. To prove this hypothesis, we conducted a long-rang RT-qPCR assay to investigate the integrity of PEDV-HB3 genome after exposing PEDV virions to the E-beam irradiation. Given that photons or electrons might cause random breaks or damage to viral RNA strand during E-beam irradiation, we designed three sets of qPCR primers, which were located at 746 bp, 3655 bp and 7674 bp upstream of the reverse transcription (RT) priming site respectively, to evaluate the integrity of viral genome (Figure 4A, Table 2). The target site of primer set 1 was located in the S gene which encodes the spike protein. The target sites of primer set 2 and primer set 3 were located in ORF1a/b that encodes the replicase (Figure 4A). As shown in Figure 4B-D, all three target sites for evaluation showed that E-beam irradiation damaged the viral RNA in a dose-dependent manner. Of note, for the long-range RT-qPCR targeted site that was 746 bp from RT priming site, E-beam irradiation with 5 kGy dose did not show significant decrease on intact RNA abundance compared to the non-treatment control group (Figure 4B). This might be due to the target region was too close to the RT priming site and the chance of viral RNA genome damage caused by 5 kGy radiation dose in the 746 bp-long region remains low. The target region that located at longer distance from RT priming site would be conducive to the accuracy of evaluation since the RT product is over 8000 bp. For the target site 3655bp from the RT priming site, 5 kGy radiation dose led to a significant decrease on the intact RNA abundance (Figure 4C). As shown in Figure 4D, E-beam irradiation with 5kGy, 10 kGy, 20 kGy, 25 kGy and 30 kGy dose decreased intact RNA abundance to 46.25%, 62.92%, 86.63%, 92.11% and 90.00% respectively. Combining together, we inferred that genome damage might be the root cause for E-beam-induced PEDV inactivation.

**Figure 4.**
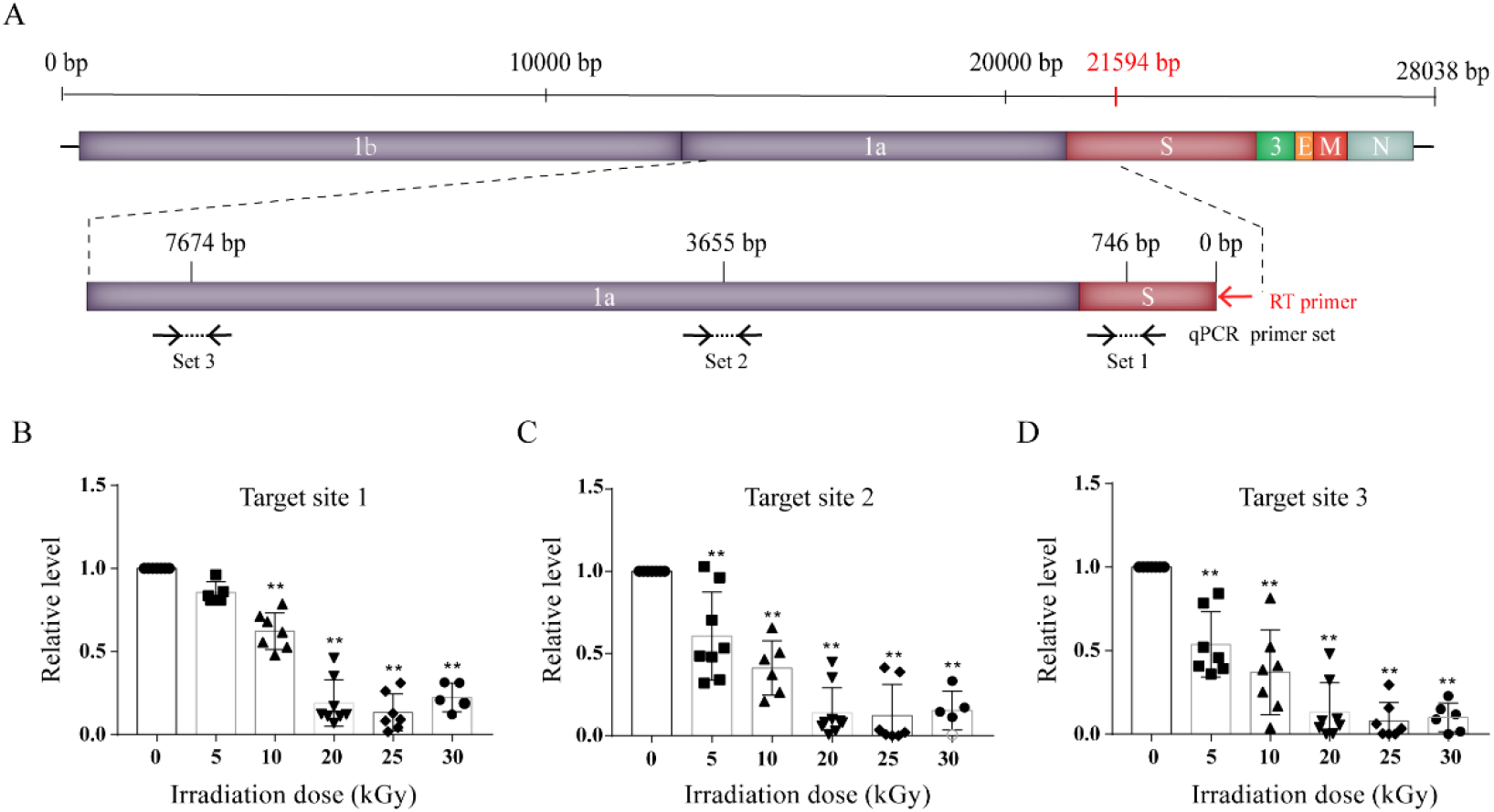
The effect of E-beam irradiation on PEDV genome. (A) Schematic diagram of PEDV-HB3 genome, RT primer site and three pairs of qPCR primer sites. Specific RT primer was indicated by red arrow. (B)(C)(D) Long-range RT-qPCR quantitative results of three target sites. The relative RNA abundance response to no irradiation is plotted. Error bars represent the standard deviation (SD) of replicate experiments. Significance was determined using a one-way analysis of variance (ANOVA) followed by Dunnett’s multiple comparisons test (p<0.05). 1a/b: ORF1a/b, replicase; S: spike protein; 3: ORF3; E: envelope protein; M: membrane protein; N: nucleocapsid protein.

## 4. Discussion

Evidence reports that the nuclei acids of SARS-CoV-2 have been isolated from cold chain frozen food and package surface, and even viable SARS-CoV-2 were detected on the cold chain outer package surface, which makes the cold chain a potential pathway for the spread of the virus (9, 25). It is well known that the low temperature increases the stability of SARS-CoV-2(26). Our study also demonstrated PEDV, a surrogate of SARS-CoV-2, remained viable at least 30 days at -20 °C. Moreover, the human immune system would be less effective at low temperatures, the SARS-CoV-2 infects relevant workers through cold chain transportation remains possible. So far, heating, chemical disinfection and UV irradiation are the mainstream methods to inactivate SARS-CoV-2. However, the usage of these methods is limited in the cold chain disinfection due to the low temperatures. This study suggested that E-beam irritation is an efficient method to inactivate coronavirus in cold chain frozen condition.

At -20 °C, E-beam irradiation with 10 kGy dose deactivated the PEDV on the surface of corrugated cardboard, polystyrene plastics and polystyrene foam. Notably, 10 kGy E-beam can indiscriminately inactivate PEDV on both the top and bottom surfaces of 18 mm thick polystyrene foam. The Food and Agriculture Organization of the United Nations (FAO), the World Health Organization (WHO), and the International Atomic Energy Agency (IAEA) stated in the 1980s that “any food has no toxicological hazard when its overall average absorbed dose does not exceed 10 kGy, no longer required to do toxicology test, and it is also safe in nutrition and microbiology.” Moreover, this study showed that a radiation dose of 5 kGy abolished 96.3% of virus infectious titers. In addition, the possible titer of virus in real cold chain transportation was much lower than that used our study. Therefore, we infer that the dose of the E-beam to inactivate coronavirus would be between 5 kGy and 10 kGy in actual application in cold chain industry. Compared to gamma irradiation, E-beam requires shorter irradiation time, processes with more precise dosage with lower variances and is more environment-friendly (27). In conclusion, E-beams is a preferable candidate for the safe and green application to inactivate SARS-CoV-2 in the cold chain.

Regarding the mechanism of E-beam inactivating viruses, it has been proposed that the E-beam preferentially destroys viral nucleic acids to inactivate Tulane virus (TV), murine norovirus (MNV-1) and Influenza A (H3N8) (17, 18). E-beam irradiation damage viral nucleic acids directly and indirectly. Direct damage means that the electron beam can directly attack the viral nucleic acid, resulting in the breakage of the viral nucleic acid sugar-phosphate backbone or the deletion, substitution and insertion of viral nucleic acid bases(28, 29). For the indirect damage, the electron beam first reacts with H_2_O to form various active particles such as hydroxyl radicals, hydrated electrons and hydrogen. These active particles then react with viral nucleic acid to cause nucleic acid fragmentation and cross-linking(30, 31). The break in the nucleic acid strand is generally sufficient to inactivate the virus (32). Consistent with this, the copy number of integrated PEDV genome decreased with increasing radiation dose. Our data implied that viral genomic RNA damage might be the main mechanism of E-beam-induced PEDV inactivation.

Previous studies have reported that low-energy electron beams and γ -ray irradiation do not affect the structure of viral proteins. In our experiments, we found that the integrity of the protein capsid was disrupted by E-beam irradiation with 20 kGy or higher dose. We speculated that this discrepancy is due to the radiation dose difference. Predmore et al. also reported that high dose of high-energy electron beam could cause damage to the Murine norovirus 1 (MNV-1) capsid (17). Since the destruction of the protein capsid and lipid bilayer envelope can further lead to a reduction in the pathogenicity of virus (33-35), a little bit higher dose of E-beam irradiation would be applied for a tough scenario to disinfect SARS-CoV-2 in cold chain.

Taken together, this study provides useful baseline data toward a potential efficient and green technique to inactivate the SARS-CoV-2 virus in cold chain environment. In the future, it will be helpful to develop more practical E-beam instruments to disinfect the SARS-CoV-2 virus in cold chain industry and prevent the cold chain as vectors for the transmission of COVID-19.

## Acknowledgements

We are grateful to Dr. George F. Gao for kindly providing Vero E6 cell. We acknowledge Dr. Jinghua Yan for kindly providing PEDV-HB3 strain. We thank Fulian Liao, Yonglei Yang for helpful suggestions and comments, Yaxin Zhu for BLS-2 facility. This research was supported by the National Natural Science Foundation of China Grant (52091541), National Key R&D Program of China Grant (2019YFA0905500), Chinese Academy of Sciences Grant (GQRC-19-18), and the Senior User Project of RV KEXUE (KEXUE2019GZ05).

## Notes

### Competing Interest Statement

The authors have declared no competing interest.

